# Inhibition of classical and alternative modes of respiration in *C. albicans* leads to cell wall remodelling and increased macrophage recognition

**DOI:** 10.1101/330167

**Authors:** Lucian Duvenage, Louise A. Walker, Aleksandra Bojarczuk, Simon A. Johnston, Donna M. MacCallum, Carol A. Munro, Campbell W. Gourlay

## Abstract

The human fungal pathogen *C. albicans* requires respiratory function for normal growth, morphogenesis and virulence. As such the mitochondria represent an enticing target for the development of new antifungal strategies. This possibility is further bolstered by the presence of fungal specific characteristics. However, respiration in *C. albicans*, as is the case in many fungal organisms, is facilitated by redundant electron transport mechanisms that makes direct inhibition a challenge. In addition, many chemicals known to target the electron transport chain are highly toxic. Here we make use of chemicals with low toxicity in mammals to efficiently inhibit respiration in *C. albicans*. We find that use of the Nitric Oxide donor, Sodium Nitroprusside (SNP), and the alternative oxidase inhibitor, SHAM, prevent respiration, lead to a loss in viability and to cell wall rearrangements that increase the rate of uptake by macrophages *in vitro* and *in vivo*. We propose that SNP+SHAM treatment leads to transcriptional changes that drive cell wall re-arrangement but which also prime cells to activate transition to hyphal growth. In line with this we find that pre-treatment of *C. albicans* with SNP+SHAM leads to an increase in virulence. Our data reveals strong links between respiration, cell wall remodelling and activation of virulence factors. Our findings also demonstrate that respiration in *C. albicans* can be efficiently inhibited with chemicals which are not damaging to the mammalian host, but that we need to develop a deeper understanding of the roles of mitochondria in cellular signalling if they are to be developed successfully as a target for new antifungals.

**Author Summary:** Current approaches to tackling fungal infections are limited and new targets must be identified to protect against the emergence of resistant strains. We investigate the potential of targeting mitochondria, organelles required for energy production, growth and virulence, in the yeast human fungal pathogen *Candida albicans*. Our findings suggest that mitochondria can be targeted using drugs that can be tolerated by humans and that this treatment enhances their recognition by immune cells. However release of *C. albicans* cells from mitochondrial inhibition appears to activate a stress response that increases traits associated with virulence. Our results make it clear that mitochondria are a valid target for the development of anti-fungal strategies but that we must determine the mechanisms by which they regulate stress signalling and virulence ahead of successful therapeutic advance.

## Introduction

*C. albicans* is one of the most prevalent fungal pathogens and a major cause of nosocomial infections which have a high mortality rate [1]. Although effective, current antifungals target a limited number of cellular processes and the development of new therapeutic approaches is essential. *C. albicans* requires mitochondrial function for normal growth, morphogenesis and virulence [2–4] but mitochondria have not been exploited as a therapeutic target to date. However given the central role of this organelle in processes essential for growth, maintenance and adaptability, coupled to the presence of fungal specific characteristics, it may be possible to develop therapies based on mitochondrial inhibition.

*C. albicans* is a Crabtree Negative yeast and relies mainly on oxidative phosphorylation for ATP production during growth and morphogenesis. It possesses a classical electron transfer chain (ETC), consisting of Complexes I-IV, as well as a cyanide-insensitive alternative oxidase, which permits respiration when the classical chain is inhibited (Fig 1A) [5]. A functional electron transport system has been shown to be important for aspects of *C. albicans* biology that are linked to virulence. For example, the inhibition of respiration in *C. albicans* and other pathogenic fungi leads to decreased growth rate [6]. Mutants defective in respiration have consistently been shown to affect the hyphal morphological switch, an important determinant of virulence in *C. albicans*, whether negatively (*goa1*Δ, *ndh51*Δ) [7] or positively (*dbp4*Δ, *hfl1*Δ, *rbf1*Δ) [8]. Although it remains unclear how mitochondria interface with the control of hyphal transition, signalling between complex II and Co Q has been shown to play a role [9].

**Fig 1.**
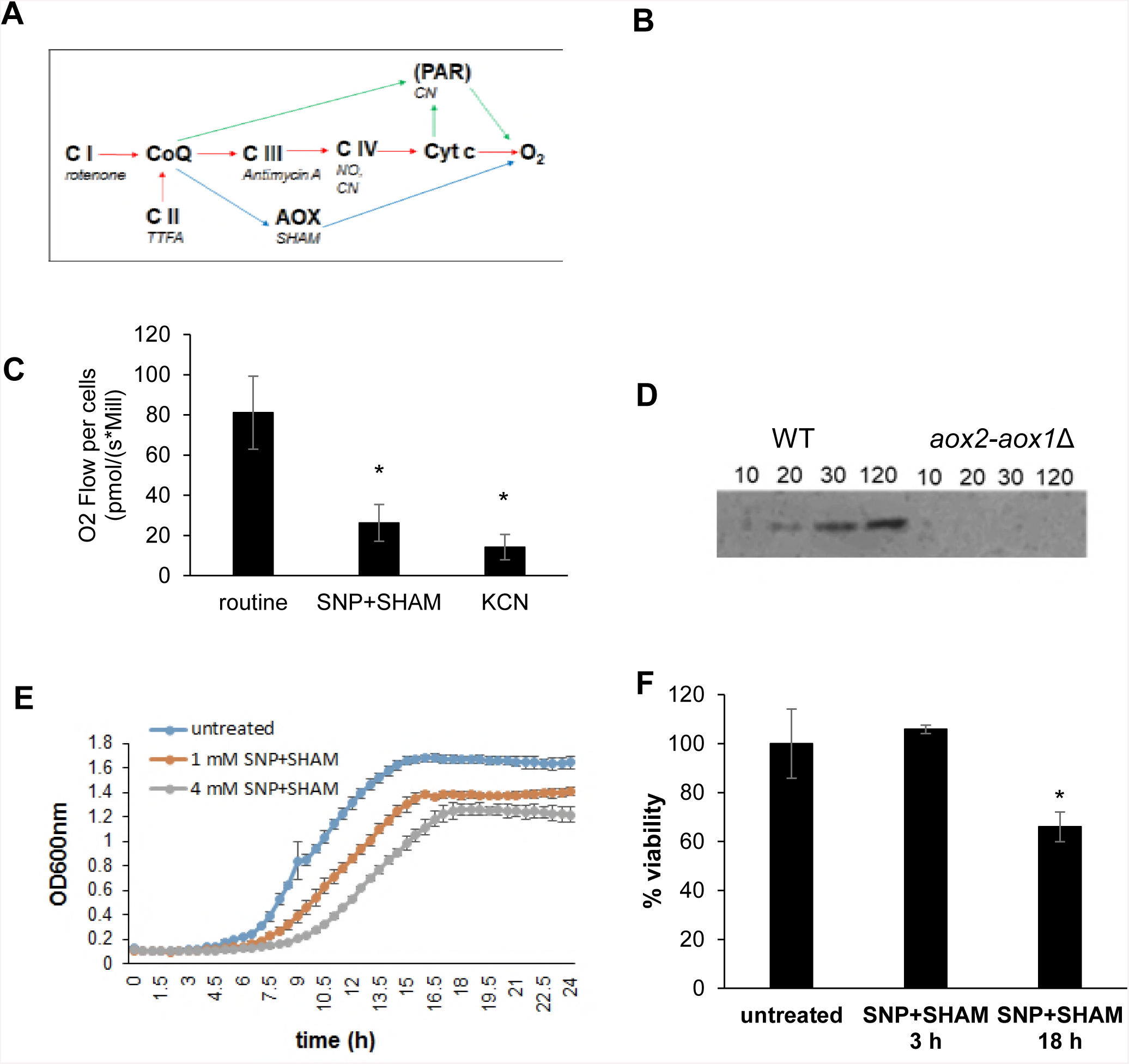
SNP+SHAM inhibits respiration and reduces viability. **(A)** Schematic of electron transport pathways in *C. albicans*, representing complexes I-IV of the classical pathway (red arrows), the alternative oxidase pathway (AOX) (blue arrows) and the parallel pathway (PAR: not shown in detail) (green arrows). Shown in italics are commonly used inhibitors. (**B)** A representative example of respiration in whole *C. albicans* cells determined using high-resolution respirometry. SNP and SHAM were added where indicated resulting in final concentrations of 1-and 2 mM for both. Potassium cyanide (KCN) was added to a final concentration of 2 mM. (**C)** Respiration was inhibited by SNP+SHAM or 2 mM KCN and compared to untreated controls, n=6. *p<0.01. (**D)** Aox2 expression was monitored by immunoblotting following addition of 1 mM SNP in samples taken from *aox2*-*aox1*Δ or control cells. (**E)** Growth was monitored over time in the presence of increasing concentrations of SNP+SHAM. (**F)** Cells were grown in the presence of 1 mM SNP + 0.5 mM SHAM for 3 h and 18 h before determining viability by colony forming unit assay, n=4. Graphs show means ± standard deviation. Student’s t-test was used to compare groups, *p<0.01

The cell wall is critical for viability of fungal pathogens and is a major determinant of virulence. Its composition and organisation also determines the recognition of *C. albicans* by the immune system [10–12]. Recent work has shown that masking of cell wall components facilitates immune evasion. Changes in surface beta-glucan exposure can occur in response to a variety of stimuli including change in carbon source and pH [13,14]. These findings are important as the majority of existing antifungals target the cell wall either directly or indirectly through the inhibition of ergosterol synthesis [15]. Changes in cell wall structure can therefore influence the outcome of *C. albicans* infections. A number of studies suggest that mitochondrial function may be linked to the maintenance of the *C. albicans* cell wall. Loss of the Complex I regulator, Goa1, revealed a link between respiration and sensitivity to cell wall damaging agents [16] and cell wall architecture [17]. In addition, impairment of mitochondrial function by deletion of *SAM37*, which encodes a component of the mitochondrial outer membrane Sorting and Assembly Machinery (SAM) complex also resulted in defects in cell wall integrity [18]. A better understanding of the links between mitochondrial function and virulence and indeed the consequences of respiration inhibition are crucial if we are to develop mitochondria as potential drug target.

In an attempt to understand the consequences of respiration inhibition in *C. albicans* we adopted a pharmacological approach. Complex I can be inhibited by Rotenone, Complex III by Antimycin A and complex IV by Cyanide (Fig 1A). However, although effective as research tools, these compounds are highly toxic and unsuitable for use in humans. As an alternative to cyanide compounds we can make use of nitric oxide (NO) which inhibits respiration at the level of Complex IV and has a safe record for use as a vasodilator [19] and in dermatologic applications [20]. NO is a molecule produced by phagocytes as an antimicrobial and is known to inhibit respiration in bacteria and fungi. NO has been evaluated as therapy in a variety of bacterial and fungal infections, including *Pseudomonas aeruginosa* in cases of cystic fibrosis and infections caused by dermatophytes [21–23]. NO works by the inhibition of Cytochrome c Oxidase at low concentrations but is rapidly reversible by oxygen. However permanent inhibition of respiration can result at higher NO concentrations [24]. In addition, NO causes the formation of reactive nitrogen species such as peroxynitrite which can damage mitochondrial function and have been shown to have strong antifungal activity [25]. Several studies report the efficacy of NO against *C. albicans* [26–28]. NO nanoparticles or NO-embedded medical devices have an anti-biofilm effect on *C. albicans*, and NO has also been shown to be effective as an adjunctive therapy in combination with existing drugs [29]. The alternative oxidase can be inhibited by hydroxamic acids such as salicylhydoxamic acid (SHAM). The low toxicity *in vivo* of alternative oxidase inhibitors such as SHAM and ascofuranone has been evaluated in their ability to treat trypanosomiasis [30,31].

We find that *C. albicans* cells are highly adaptive to classical respiration inhibition but that a combination of SHAM and the NO donor Sodium Nitroprusside (SNP) leads to fitness defects and loss of viability. In addition, SNP+SHAM treatment leads to cell wall organisation defects that unmask *C. albicans* cells leading to increased immune cell recognition in cell culture and animal models. However, release of cells from SNP+SHAM treatment leads to a rapid activation of the hyphal transition programme and increased virulence in a mouse model. Our data suggest that mitochondria form part of a complex response network that is imbedded within cellular responses that are important for *C. albicans* virulence.

## Results

### SNP+SHAM inhibits respiration, growth and reduces viability in *C. albicans*

Our goal was to identify conditions that would allow for the reproducible inhibition of classical and alternative respiration in *C. albicans*. To achieve this oxygen consumption was measured upon exposure of cells to the nitric oxide donor SNP which inhibits cytochrome c oxidase, and the alternative oxidase (Aox) inhibitor SHAM. Initial titration experiments suggested that the concentrations of 1 mM SNP and 0.5 mM SHAM were suitable for our purpose. Recovery from NO induced inhibition of classical respiration by SNP addition was rapid, with full restoration achieved within 30 min (Fig 1B). Further application of SNP did not lead to a reduction in oxygen consumption suggesting that cells had switched to alternative respiration upon SNP treatment (Fig 1B). This was confirmed by subsequent addition of SHAM, which decreased the respiration level significantly (Fig 1B). Addition of 2mM cyanide, which has been shown to inhibit the parallel pathway in *C. parapsilosis* [44] was sufficient to inhibit the remaining respiration suggesting the presence of a parallel pathway in *C. albicans*. Addition of SNP+SHAM simultaneously led to significant loss of respiration that was comparable to that observed upon cyanide addition (Fig 1C). In support of a rapid transition to alternative respiration, an increase in Aox2 protein levels was detectable within 20 min following exposure to SNP (Fig 1D).

Given the effects of SNP+SHAM on respiration we investigated whether co-treatment led to effects on growth and viability. Incubation of *C. albicans* cells with SNP+SHAM led to a decrease in growth and final biomass in a dose dependent manner (Fig 1E). Treatment with SNP+SHAM for 3 hours did not have a significant effect on viability, however, prolonged SNP+SHAM exposure resulted in decreased viability when compared to untreated controls (Fig 1F). These data suggest that *C. albicans* possesses a robust and adaptable respiratory system that can be targeted using a SNP + SHAM co-treatment approach.

### SNP+SHAM treatment induces structural re-arrangements within the cell wall

Previous studies suggest a link between respiration and cell wall integrity [8,16]. Our goal was to determine whether chemical inhibition of respiration using SNP+SHAM could negatively impact cell wall integrity. SNP+SHAM exposure led to a reduction in growth and sensitivity to the cell wall damaging agents Calcoflour White (CFW) and Congo Red (CR) (Fig 2A). We employed transmission electron microscopy to examine cell wall structure following SNP+SHAM treatment (Fig 2B). This analysis revealed that the outer wall, composed of mannans and cell wall proteins, but not the inner cell wall, was reduced in thickness when compared to untreated cells (Fig 2B and C). The density of mannan fibres in this layer was also found to be increased by SNP+SHAM treatment. These data suggest that SNP+SHAM causes cell wall changes which result in an altered organisation of the outer cell wall. HPLC analysis of acid-hydrolysed cell wall components showed that SNP+SHAM treatment caused no significant changes in the bulk levels of chitin, glucan or mannan (Fig 2D). This suggests cell wall changes induced by SNP+SHAM are due to changes in wall organisation rather than gross changes in the levels of cell wall components. Interestingly we also observed the presence of a large structure adjacent to the vacuole within SNP+SHAM treated cells (Fig 2E). We confirmed this to be a lipid droplet using the neutral lipid stain LD540 (Fig 2F). This finding suggest a link between respiratory inhibition and the regulation of lipid biosynthesis in *C. albicans*.

**Fig 2.**
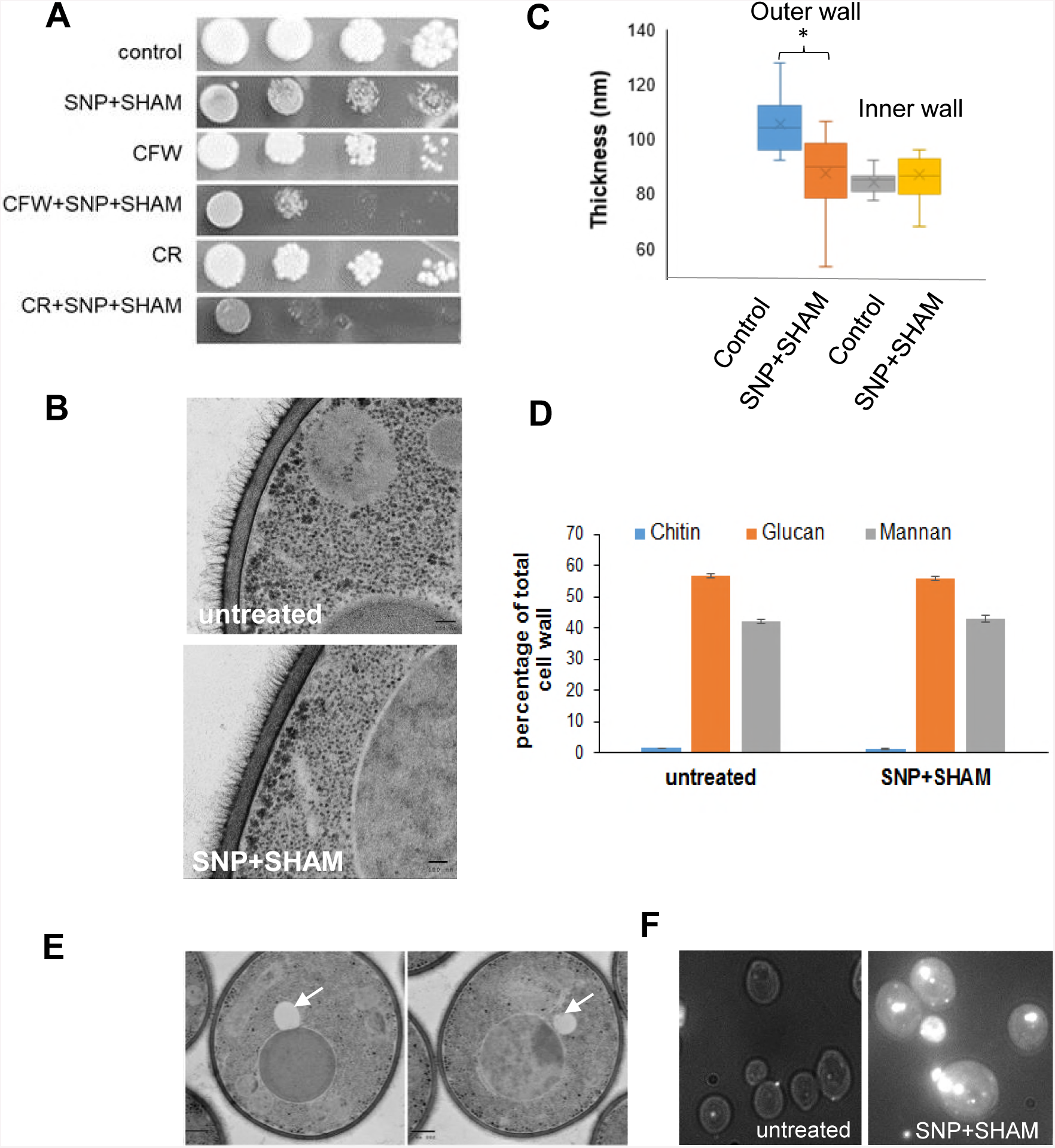
SNP+SHAM treatment leads to cell wall alterations and altered lipid metabolism. **(A)** Wild-type cells were serially diluted and equal volumes were spotted onto YPD plates containing 1 mM SNP+ 0.5 mM SHAM and 25 μg/ml Calcofluor White (CFW) or 40 μg/ml Congo Red (CR). **(B)** Cells were treated with 1 mM SNP + 0.5 mM SHAM for 18 h and processed for TEM. Representative examples of untreated and SNP+SHAM treated cells are shown, size bar = 100 nm. **(C)** Quantification of outer-and inner cell wall thicknesses was carried out from TEM images. Measurements were taken at 5 points along the cell wall, n=25. (**D)** Cell wall material was extracted and acid hydrolysed from cells treated with 1 mM SNP + 0.5 mM SHAM for 18 h, followed by HPLC analysis, n=3. Graphs show means ± standard deviation. (**E)** TEM images from cells treated with SNP+SHAM, arrows indicate lipid droplets (**F)** Control and SNP+SHAM treated cells were stained using the neutral lipid stain LD540 and viewed by fluorescence microscopy, size bar = 10 μm. Student’s t-test was used to compare groups, *p<0.01.

### Caspofungin resistance reveals a link between respiration inhibition and Upc2

Cell wall damage can be sensed and repaired by activation of the cell wall integrity pathway (CWI) [45]. To determine whether this mechanism was engaged by SNP+SHAM treatment CWI pathway activity was measured by detection of the phosphorylated forms of Mkc1, a well characterised marker of pathway activation [46]. We also determined whether SNP+SHAM treatment led to the activation of Hog1, a major oxidative stress response regulator that also influences cell wall biosynthesis [47]. Unlike the positive controls, SNP+SHAM treatment did not cause activation of Hog1 or Mkc1 (Fig 3A). This suggests that either the major cell wall integrity responses are not involved in SNP+SHAM-induced cell wall changes or that the mitochondrial compartment functions downstream of Hog1 and Mkc1 within the cell wall response.

**Fig 3.**
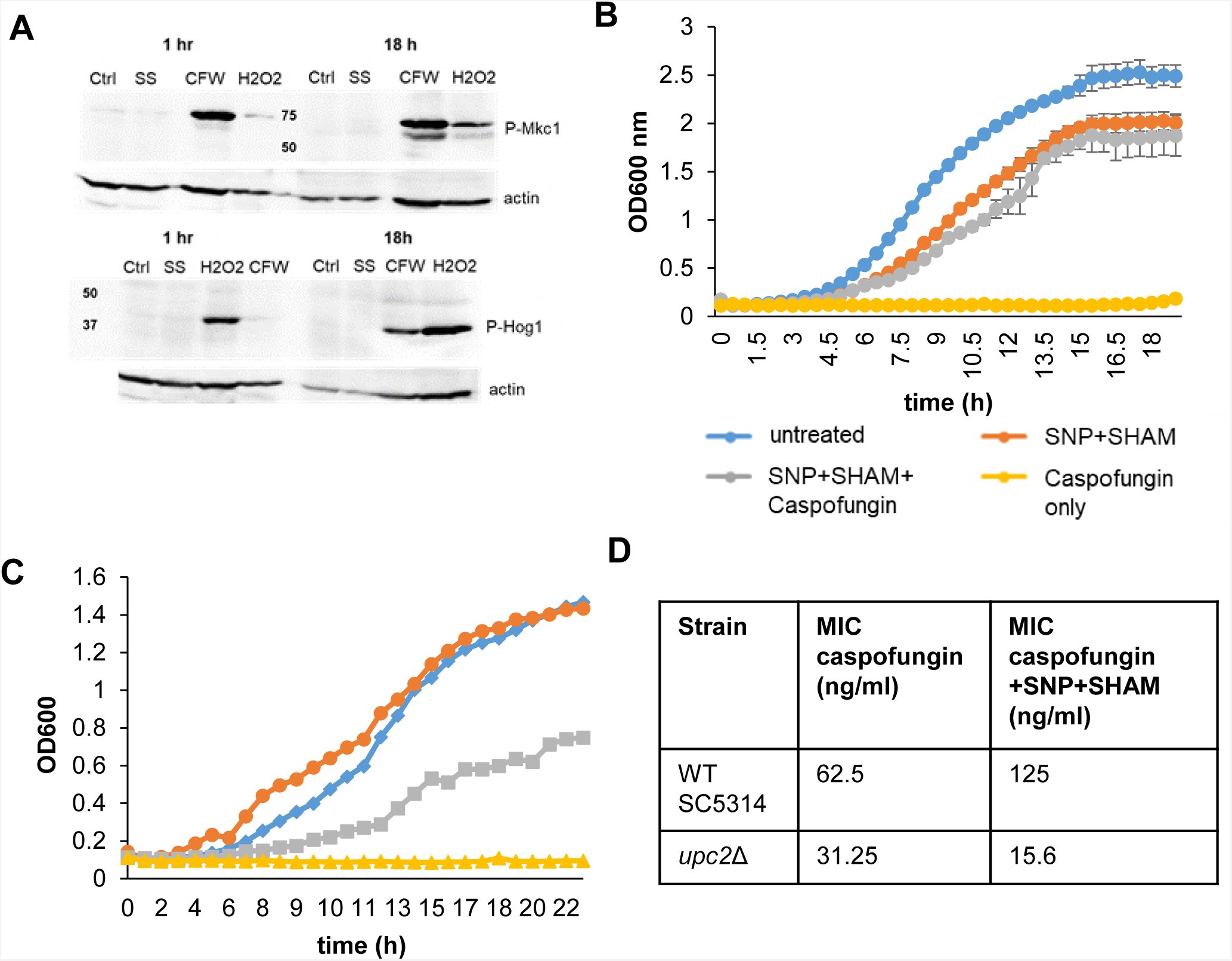
SNP+SHAM-induced cell wall changes does not induce Mkc1 or Hog1 activation but alters caspofungin sensitivity, which requires Upc2. **(A)** Cells were treated with 1 mM SNP + 0.5 mM SHAM (SS) for 1 h or 18 h. Activation of Mkc1 was determined using a p44/42 antibody. Cells treated with 25 μg/ml Calcofluor White (CFW) were used as a positive control. Activation of Hog1 was determined using a p38 antibody, using cells treated with 2 mM hydrogen peroxide as a positive control. The level of α-actin on the respective blots is shown as a loading control. (**B)** Wild-type cells were grown with 1 mM SNP + 0.5 mM SHAM and/or 100 ng/ml caspofungin, n=3. (**C**) *upc2*Δ and its respective wild type were grown in synthetic complete medium with 1 mM SNP+ 0.5 SHAM or in combination with 100 ng/ml caspofungin. (**D)** Microdilution assays with 3.9 ng/ml – 8 μg/ml caspofungin with or without 1 mM SNP + 0.5 mM SHAM were performed. The table shows the MICs for caspofungin for the wild-type strain and the *upc2*Δ mutant with or without SNP+SHAM, n=3.

To identify potential regulators of the SNP+SHAM response that may be important in cell wall organisation we treated *C. albicans* with a combination of SNP+SHAM and caspofungin, which targets the cell wall (Fig 3B). We observed that SNP+ SHAM treatment led to caspofungin resistance, presumably a result of the alteration in cell wall structure (Fig 3B). Using this phenotype, we screened a library of transcription factor deletion mutants [32] by growing the strains in the presence of SNP+SHAM in combination with caspofungin at a level that inhibited growth of the wild type strain. A single transcription factor, Upc2, was found to be required for caspofungin resistance upon SNP+SHAM treatment. Deletion of *UPC2*, which plays an important role in the regulation of ergosterol biosynthesis, led to a reproducible reduction in caspofungin resistance upon exposure to co-treatment with SNP+SHAM (Fig 3C). To confirm the results from the initial screen, microdilution growth assays of caspofungin in combination with SNP+SHAM were performed. The results showed that the presence of SNP+SHAM increased the MIC of caspofungin against the wild type strain SC5314 (Fig 3D). An independently-derived *upc2*Δ mutant [48] was slightly more sensitive to caspofungin, nevertheless, SNP+SHAM decreased the caspofungin MIC instead of increasing it as in the wild-type strain (Fig 3D). These data may suggest a role for Upc2 in regulating cell wall changes in response to SNP+SHAM treatment.

### Analysis of transcriptional changes in response to SNP+SHAM treatment

To further examine the changes caused by SNP+SHAM we examined the transcriptional response of *C. albicans* to this treatment using RNAseq. RNA was extracted from log-phase yeast cells exposed to 1 mM SNP, 0.5 mM SHAM, or both for 30 minutes (Fig 4A and B and Supplementary Tables S2-S4). A short exposure time was selected in order to capture early alterations as opposed to profiles from the expansion of adapted cell lineages. SNP and SNP+SHAM treatment led to the differential expression of over 1500 genes, whereas SHAM treatment led to a smaller response with only 131 genes differentially expressed (Fig 4A and supplementary tables S2-S4). A significant overlap between differentially expressed genes within SNP and SNP+SHAM treatment data sets was observed (Fig 4A). As expected the alternative oxidases *AOX2* and *AOX1* and genes required for a response to nitric oxide, such as *YHB1*, were upregulated in both SNP only and SNP+SHAM groups. Several components of classical respiration were downregulated, upon SNP+ SHAM treatment while genes involved in glyoxylate cycle (*MLS1* and *ICL3*) and glycolysis and gluconeogenesis (*FBP1, GPM1, GPM2, PCK1, PGI1, PGK1*) (Supplementary Tables S2 and S4) were upregulated. These changes suggest a shift in carbon metabolism in response to respiration inhibition.

**Fig 4.**
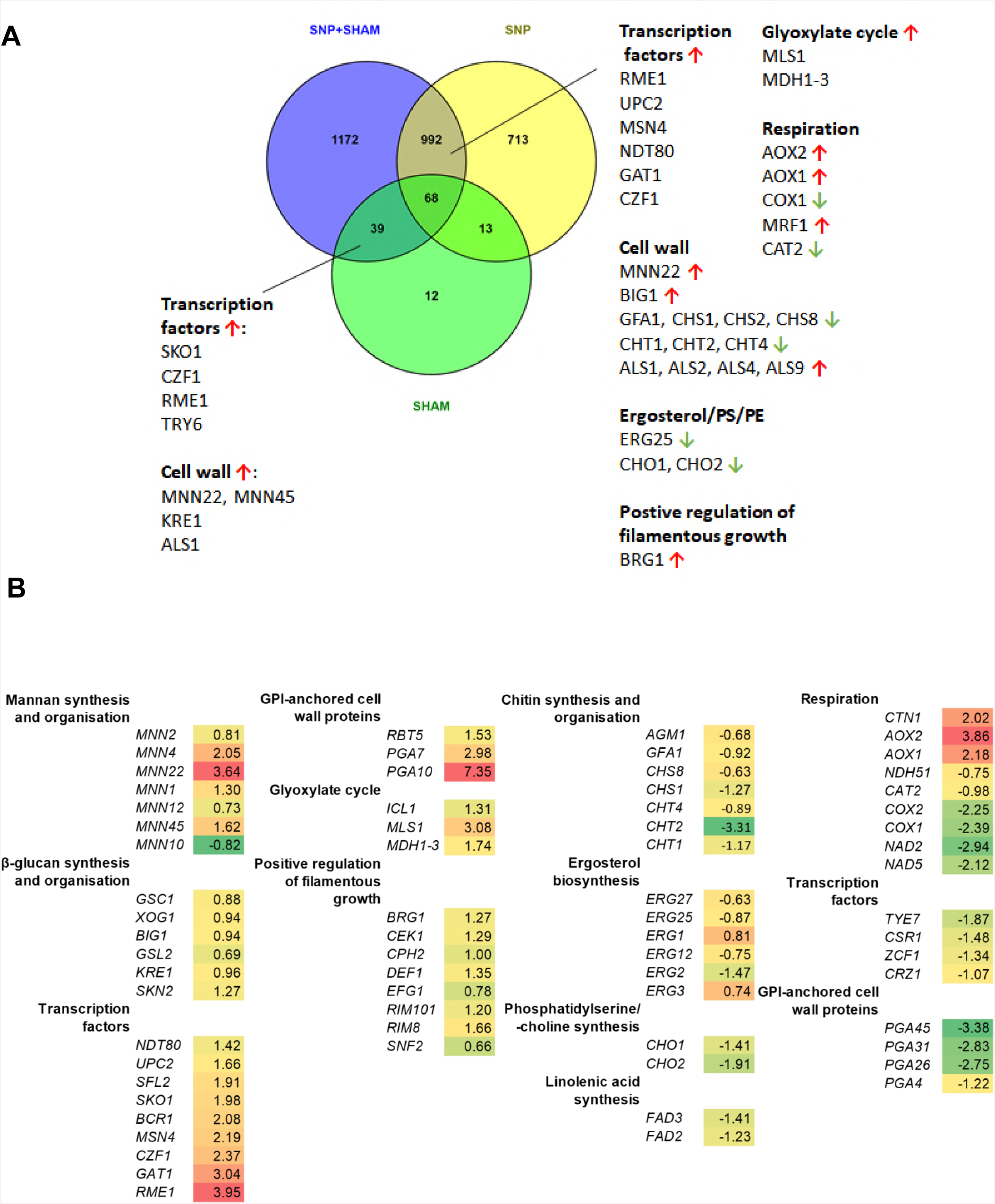
Transcriptome of wild-type *C. albicans* treated with SNP, SHAM or SNP+SHAM. (**A)** Venn diagram showing overlap in differentially expressed genes between SNP, SHAM and SNP+SHAM treatments by RNAseq analysis as described in materials and methods. (**B)** A selection of differentially expressed genes in SNP+SHAM treated cells were grouped by GO term, with log_2_-fold change vs. untreated shown. Green depicts downregulated genes and red/yellow depicts upregulated genes.

A number of transcriptional regulators were upregulated in response to both classical and alternative respiration inhibition (Fig 4A). To examine which key regulators were involved in the response to SNP+SHAM we analysed the differentially expressed gene list using PathoYeastract, a tool which ranks transcription factors in order of the number of genes in the list that they regulate, for which there exists supporting expression evidence [49]. This analysis showed that that the transcription factor Sko1 has regulatory associations with 33.8 % of the differentially expressed genes. *SKO1* itself was upregulated upon SNP+SHAM treatment. Interestingly, the upregulation of *SKO1* seems to be a SHAM-specific effect, as it was not differentially expressed in the SNP only dataset. Upc2 binding sites were identified within the promoters of 13.4% genes differentially regulated upon SNP+SHAM treatment. This along with the fact that *UPC2* expression was itself upregulated upon SNP+SHAM treatment further suggests a role for this transcription factor.

Analysis of SNP+SHAM RNAseq data showed that several genes involved in chitin synthesis and organisation were downregulated, including the chitinases *CHT1, CHT2* and *CHT4* (Fig 4B). Mannan biosynthesis and organisation genes were upregulated, as were genes involved in β-glucan synthesis and organisation. As no significant changes in relative glucan, mannan or chitin levels were detected by HPLC in acid-hydrolysed cell walls our data may suggest that SNP+SHAM induced transcriptional changes lead to alterations in cell wall organisation as opposed to bulk synthesis. Supporting this hypothesis we observed that several GPI-anchored cell wall protein genes that play a role in crosslinking cell wall components were also differentially expressed (Fig 4B).

Genes encoding the chitinase Cht2 and GPI-anchored cell wall protein Pga26, previously shown to have roles in unmasking of cell wall β-glucan, were downregulated [14]. This appears to be a result of inhibition of classical respiration as SNP treatment alone led to reduced expression of *CHT2* and *PGA26*. The majority of genes in the so-called “Common in fungal extracellular membranes” (CFEM) family, including *PGA7, PGA10, RBT5*, and *CSA2* were upregulated [50]. These genes encode proteins with roles in iron acquisition and adhesion. The expression of several of these genes is controlled by Bcr1, a transcription factor known to be important in the hypoxic response [51] and *BCR1* was itself upregulated. Genes involved in adhesion, including several of the *ALS* genes, as well as the regulator *RIM101* were also upregulated (Fig 4B), however no significant difference was found in *in vitro* adhesion of SNP+SHAM treated cells (Supplementary Fig S1).

SNP+SHAM treatment also led to changes in genes involved in lipid metabolism. Upregulation of several lipase genes suggested a mobilisation of lipid stores. This is in agreement with our observation of increased formation of lipid droplets upon SNP+SHAM treatment (Figs 2E and 2F). A number of genes involved in the ergosterol biosynthesis pathway were downregulated (Fig 4B). In addition, genes involved in the synthesis of phosphatidylserine (PS) and phosphatidylcholine (PC) (*CHO1* and *CHO2*, respectively) and linoleic acid (*FAD3* and *FAD2*), which are important components of the cell membrane, were also downregulated. Collectively these data suggest that SNP+SHAM leads to changes in the cells lipid biosynthesis programme.

### SNP+SHAM treatment increases surface chitin and β-glucan exposure

Our data suggests that inhibition of respiration leads to cell wall rearrangement (Figs 2B-D). To investigate this further we assessed the exposure of chitin and β-glucan at the cell surface using fluorescent probes. Wheatgerm agglutinin (WGA), binds chitin in fungal cell walls, but due to its large size cannot penetrate the cell wall to stain chitin in the innermost layers where it is usually found [52]. WGA therefore typically stains structures such as bud scars where chitin is exposed, and staining of the lateral cell wall indicates that chitin is exposed at the surface of the cell wall. SNP+SHAM treated cells exhibited a greater lateral wall staining compared to untreated cells (Fig 5A, B). These results suggest that SNP+SHAM treatment causes exposure of the normally hidden chitin in the cell wall.

**Fig 5.**
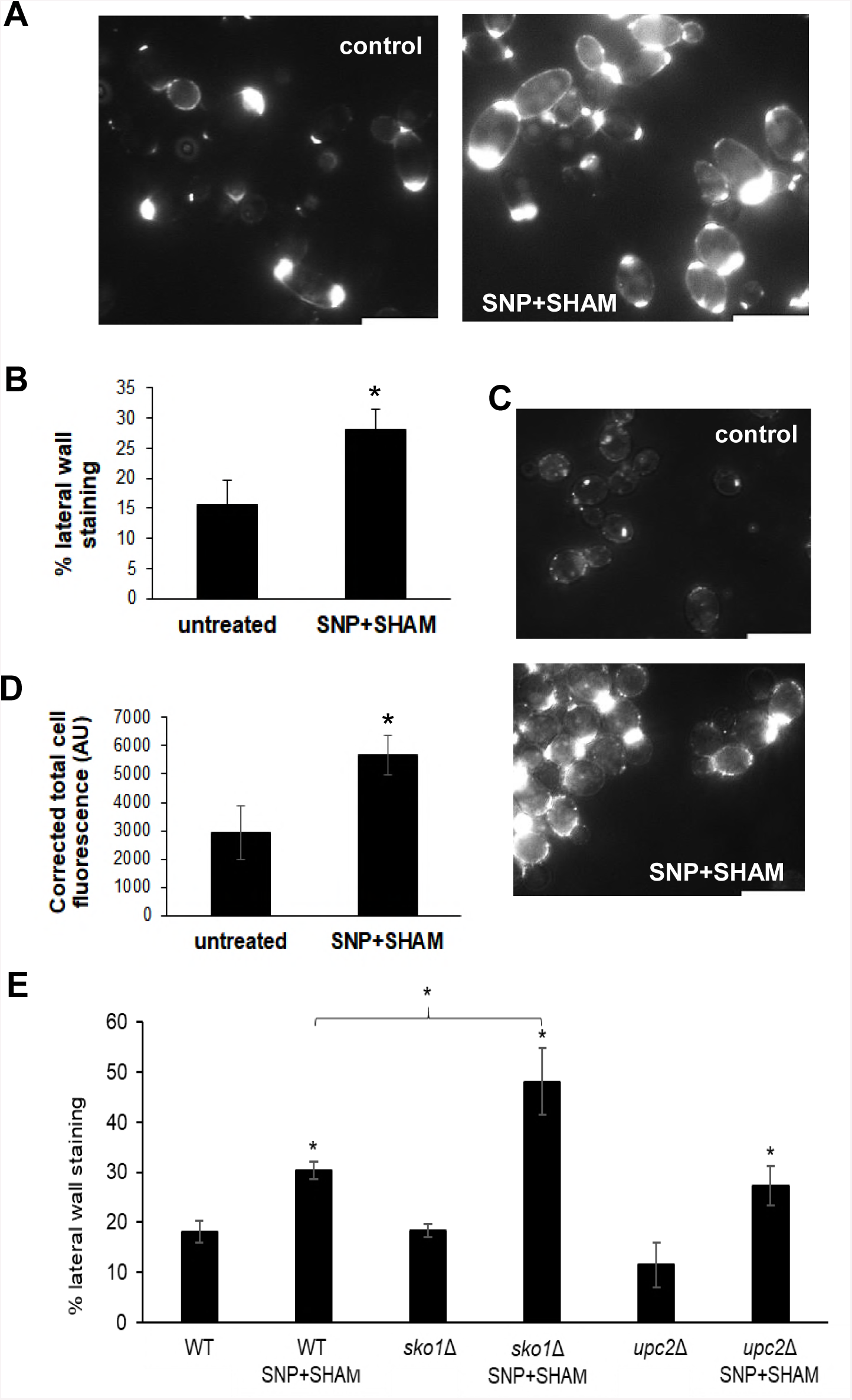
SNP+SHAM induces surface exposure of chitin and β(1, 3)-glucan. (**A)** Representative examples of untreated cells and cells treated with 1 mM SNP + 0.5 mM SHAM for 18 h stained with FITC-wheatgerm agglutinin using fluorescence microscopy and **(B)** quantified as described in materials and methods, n=3. **(C)** Representative examples of untreated and SNP+SHAM treated cells stained with dectin-1, size bar = 10 μm **(D)** quantified as described in materials and methods, n=3. **(E)** Effect of SNP+SHAM treatment on chitin unmasking as determined by WGA staining in *upc2*Δ and *sko1*Δ mutants. Graphs show the means ± standard deviation. At least three independent experiments were performed for each analysis. Student’s t-test was used to compare groups, *p<0.01, **p<0.001

Dectin-1 recognises β(1,3)-glucan in fungal cell walls and is one of the major receptors involved in the immune response against fungal pathogens [53]. β(1,3)-glucan is located in the inner cell wall, covered by an outer layer of mannoprotein fibrils, minimising its exposure to immune recognition. To determine whether SNP+SHAM treatment causes a change in β(1,3)-glucan exposure in *C. albicans*, a soluble form of dectin-1 was used to stain treated cells. SNP+SHAM pre-treatment significantly increased the level of dectin-1 staining of the cell wall (Fig 5C, D). This suggests that the cell wall changes caused by SNP+SHAM treatment causes cell wall rearrangements exposing β-glucan at the cell surface.

The observed sensitivity of *upc2*Δ cells to caspofungin in the presence of SNP+SHAM suggested that Upc2 links respiration to cell wall integrity. This led us to examine the effect of SNP+SHAM on chitin and β-glucan exposure in the *upc2*Δ mutant, to determine whether Upc2 influences unmasking. The degree of surface chitin exposure in upc2Δ, and the increase in unmasking upon SNP+SHAM treatment, was not significantly different from the wild type (Fig 5E). Similarly, the results of dectin-1 staining for *upc2Δ* was also not significantly different from the wild type strain (Supplementary Fig S2). These results suggest that cell wall unmasking and sensitivity to caspofungin in response to SNP+SHAM do not share a common mechanism.

Given that Upc2 was not linked to β(1,3)-glucan or chitin exposure we tested a number of strains deleted for transcription factors identified as upregulated within our RNAseq data in response to SNP+SHAM treatment. Our investigations did not identify deletions of any single TF that could prevent β(1,3)-glucan or chitin exposure. However deletion of *SKO1* led to an increase in chitin unmasking following SNP+SHAM, exposure relative to the wild type strain (Fig 5E). This may suggest that Sko1 acts as a repressor of chitin unmasking, or simply that it is involved in the repair of cell wall rearrangements caused by this treatment.

We wished to examine whether other forms of respiration deficiency could lead to the same cell wall unmasking phenotype. The mutant *ndh51*Δ does not possess a functional Complex I and has a low respiration level. It showed a very high degree of chitin unmasking as determined by WGA staining (Supplementary Fig S3A). Dectin-1 staining was also significantly increased, although to a lesser degree than in SNP+SHAM treated cells (Supplementary Fig S3B). These results demonstrate that inhibition of classical electron transfer chain either through Complex I or Complex IV can cause unmasking of chitin and β(1,3)-glucan.

### Pre-treatment with SNP+SHAM increases phagocytosis by macrophages

Due to the higher β(1,3)-glucan surface exposure in SNP+SHAM treated cells, it was hypothesised that these cells may be recognised and engulfed more readily by macrophages. SNP+SHAM treated cells were co-incubated with J774.1 murine macrophages and were examined after 1 h by microscopy and scored for uptake. A higher proportion of the SNP+SHAM pre-treated cells were taken up by macrophages compared to untreated cells after 1 h (Fig 6A, B). A similar increase in uptake was observed when cells were pre-treated with 1 mM KCN only (Supplementary Fig S4A) suggesting that inhibition of the classical respiratory chain is responsible for this phenotype. In agreement with this, inhibition of the alternative respiration pathway alone using SHAM or use of the *aox2-aox1*Δ mutant had no significant effect on uptake (Supplementary Fig S4A). An increase in uptake of *ndh51*Δ cells was also observed but not to the same degree as SNP+SHAM pre-treated cells (Supplementary Fig S4B).

**Fig 6.**
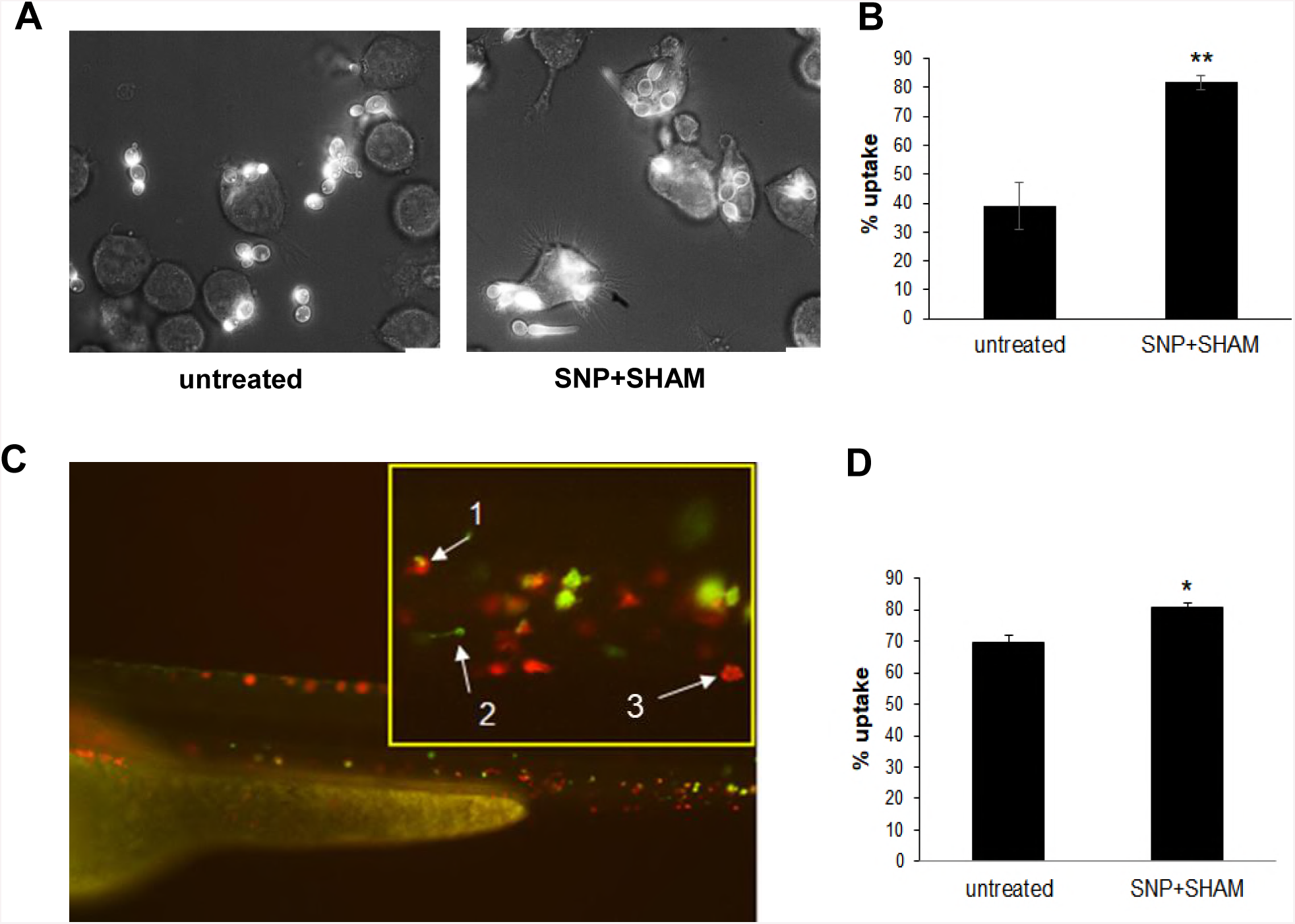
*C. albicans* pre-treated with SNP+SHAM increases uptake by murine macrophages and by zebrafish macrophages *in vivo.* **(A)** Wild-type cells were treated with 1 mM SNP + 0.5 mM SHAM for 18 h, then washed in PBS and co-incubated with J774.1 murine macrophages. Representative examples of uptake after 1 hour co-incubation are shown and (**B)** quantified by counting at least 200 *C. albicans* cells per experiment, *p<0.001, five independent experiments were carried out. (**C)** Wild-type GFP labelled *C. albicans* cells were treated with 1 mM SNP + 0.5 mM SHAM for 18 h, washed and injected into zebrafish larvae. A representative microscopy image of *C. albicans* (green) being taken up by macrophages (red) *in vivo* is presented (1 = phagocytosed *C. albicans*, 2 = single *C. albicans*, 3 = single macrophage). (**D)** Number of phagocytosed *C. albicans* cell was calculated from images taken from three independent experiments, n=3, *p<0.01, Graphs show means ± standard deviation. Student’s t-test was used to compare groups.

In order to understand how SNP+SHAM pre-treatment would influence systemic candidiasis in an infection model, we used the zebrafish larva model, in which we could examine phagocytosis of *C. albicans in vivo*. *C. albicans* expressing GFP was pre-treated with SNP+SHAM for 18h. These cells were then injected into zebrafish larvae in which macrophages were labelled with mCherry. Z-stacks of each fish were taken using confocal microscopy and uptake by macrophages was assessed manually. The uptake of pre-treated cells was approximately 10% higher than untreated cells (Fig 6C, D).

### SNP+SHAM pre-treatment increases the virulence of *C. albicans* in a mouse systemic model of candidiasis

It was not clear whether the increased uptake of SNP+SHAM pre-treated *C. albicans* by macrophages would attenuate virulence *in vivo.* To determine this, *C. albicans* were pre-treated with SNP+SHAM before being washed and injected into the tail vein of 6-8 week old female BALB/c mice and the progress of infection was monitored by weight loss and condition of the animals. Kidney fungal burden measurements were made at 3 days post-infection to give an ‘outcome score’. A higher outcome score is indicative of greater weight loss and higher fungal burdens and thus virulence. It was observed that SNP+SHAM pre-treated *C. albicans* caused a more rapid weight loss when compared to controls. Infection with SNP+SHAM treated *C. albicans* cells also led to significantly higher fungal burdens within the kidney at the time of culling. The combination of these factors led to a significantly higher outcome score for SNP+SHAM pre-treated cells compared to untreated cells (Fig 7A). This data suggests that pre-treatment with SNP+SHAM does not reduce fitness of *C. albicans* in the host, but may in fact prime a stress response that leads to increased virulence.

**Fig 7.**
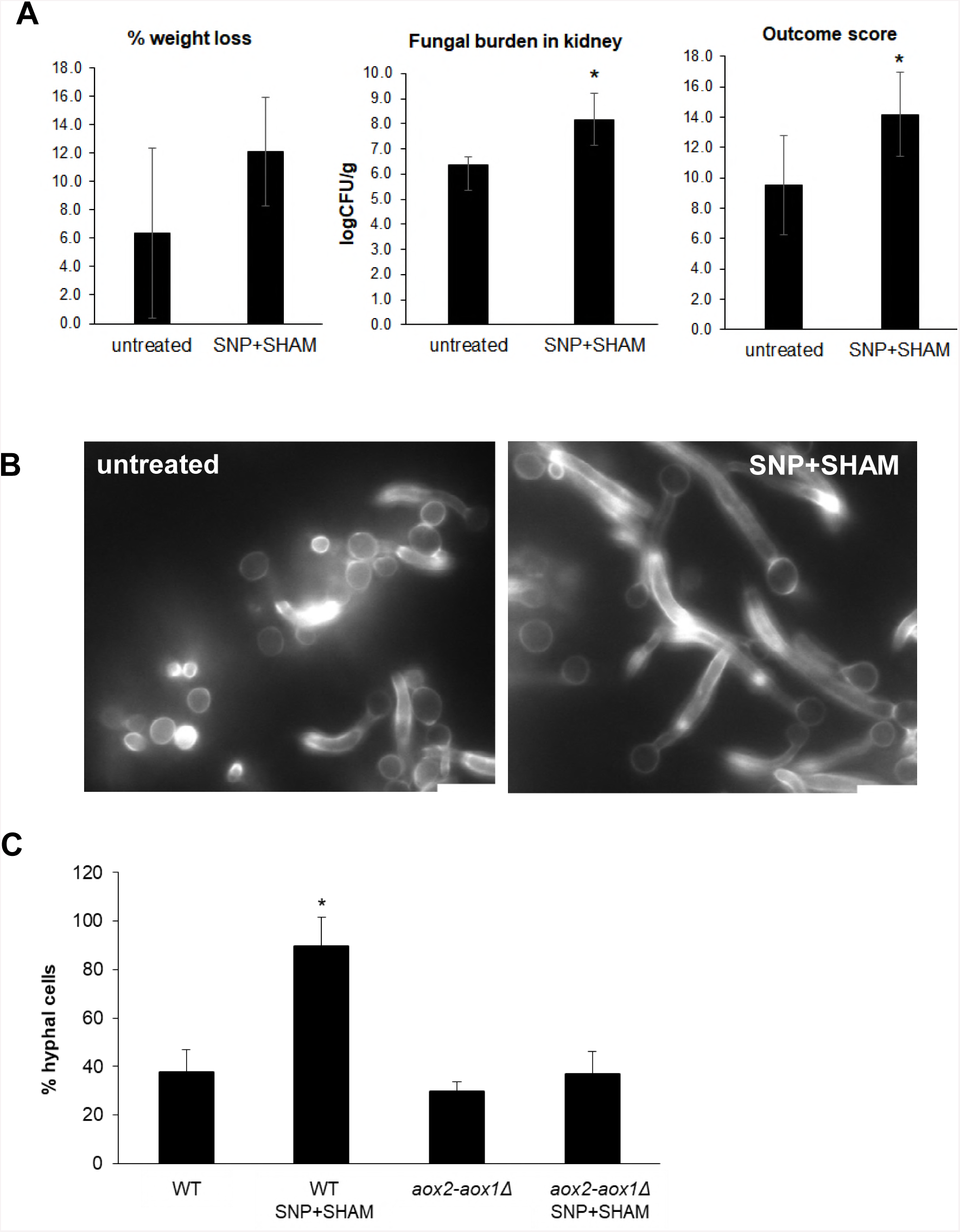
Pre-treatment of *C. albicans* increases virulence in the mouse model and leads to more rapid filamentation. **(A)** *C. albicans* cells were treated with SNP+SHAM for 18 h and then washed in PBS prior to injection via the tail vein. Weight loss, fungal burden and outcome score were calculated as described in materials and methods, *p<0.05. (**B)** Wild-type and *aox2*-*aox1*Δ mutant cells were treated with 1 mM SNP + 0.5 mM SHAM for 18 h, then washed in PBS and transferred to serum-containing medium at 37 °C. Representative examples of filamentation in wild-type untreated and SNP+SHAM pre-treated cells after 90 min. The percentage of filamentous cells was then calculated **(C),** n=3. Graphs show means ± standard deviation. Three independent experiments were performed for each analysis. Student’s t-test was used to compare groups, *p<0.01.

One possibility is that pre-treatment of *C. albicans* with SNP+SHAM may have primed cells to activate traits associated with virulence when injected into the host. We observed that cells pre-treated with SNP+SHAM exhibited a higher incidence of filamentation than untreated cells. The rapid induction of filamentation is generally associated with increased escape from macrophages and virulence [54,55]. However it was not clear whether this effect was dependent on the presence of macrophages. To determine this the incidence of hyphal formation was assessed after 90 min incubation following removal of SNP+SHAM from the medium in the absence of macrophage cells. The majority of pre-treated cells filamented within 90 min of withdrawal of SNP+SHAM, while the untreated cells were slower to form hyphae under these conditions (Fig 7B). These results suggest that SNP+SHAM treatment may prime cells for hyphal growth, which proceeds more rapidly when the inhibitors were removed. This effect was also observed when cells were pre-treated with cyanide alone (Supplementary Fig S5). Since both cyanide and NO induce alternative respiration, the ability to rapidly form hyphae after SNP+SHAM treatment was tested in an *aox2-aox1*Δ mutant. The mutant did not show the level of hyphal formation as the WT, suggesting that the alternative oxidase enzymes are required for the filamentation response upon inhibition of classical respiration.

## Discussion

Mitochondrial function has been studied extensively in the model yeast *Saccharomyces cerevisiae*, but has been a neglected area in fungal pathogens. Mitochondria have essential roles in pathogenicity [56] and antifungal drug resistance [3]. Therefore, understanding their biology is essential if we are to obtain a full understanding of fungal pathogenicity and evaluate the potential to target this organelle effectively. The electron transport chain of *C. albicans*, consisting of an alternative and parallel pathway in addition to the classical ETC [57], is highly flexible in that it allows electron flow to be re-routed should one of the pathways become blocked. Nevertheless, simultaneous inhibition of the classical and alternative pathways can inhibit respiration by up to 90 % and severely restricts growth [58].

To our knowledge, the effect of NO on respiration in *C. albicans* has not been characterised. We show NO exposure as a result of SNP treatment is effective in inhibiting respiration and activation of an alternative respiration response as has been shown for other inhibitors of classical respiration [58]. Inhibition of both classical and alternative respiration using SNP+SHAM led to sustained inhibition of respiration suggesting that the level of NO released by SNP in our experiments was sufficient to irreversibly inhibit Cytochrome c Oxidase. Peroxynitrite, which can be formed when NO reacts with endogenous superoxide, has also been shown to cause irreversible inhibition of cytochrome c oxidase [59].

SNP+SHAM showed a synergistic effect with cell wall damaging agents, Calcofluor White and Congo Red. *C. albicans* respiration-deficient mutants *dpb4*Δ, *hfl1*Δ, and *rbf1*Δ [8] and *goa1*Δ and *ndh51*Δ [16] have similarly been shown to be sensitive to CFW and CR. A number of genes found to be differentially expressed upon SNP+SHAM treatment have been reported to be involved in the response to caspofungin. Genes induced in this response include *AGP2*, *KRE1, PGA13, RHR2* and *CZF1* while repressed genes include *CHT2, PGA4* and *SCW11, HAS1, CRZ1* and *ENG1* [60–62]. These expression changes may help to explain why SNP+SHAM treated cells were more resistant to caspofungin.

Growth in the presence of SNP+SHAM caused an increase in the surface exposure of chitin and β(1,3)-glucan. This phenomenon of “unmasking” cell wall components was first described as a consequence of sub-MIC caspofungin treatment [63,64], which correlated with an increased inflammatory response by immune cells. Sherrington et al. showed that unmasking of β(1,3)-glucan and chitin occurred during adaptation to low pH [14]. The authors found no activation of Mkc1 kinase involved in the cell wall integrity response, and increased uptake by macrophages, both of which were also seen in SNP+SHAM-induced unmasking. Reduced expression of *CHT2* has been shown to enhance chitin exposure at the cell surface. *CHT2* was found to be downregulated in SNP+SHAM treated cells, suggesting a possible link to chitin exposure. Masking of β(1,3)-glucan has been shown to occur during growth on lactate, which aids in immune evasion. [13]. It was found that Crz1 was required for β(1,3)-glucan masking, and that *pga26*Δ and *cht2*Δ mutants showed elevated β(1,3)-glucan exposure. These and other genes encoding cell wall proteins were found to be downregulated in SNP+SHAM treated cells, suggesting repression of glucan masking at the level of Crz1 but also a shift in the cell wall protein profile promoting elevated β(1,3)-glucan exposure (Fig 4B).

Genes involved in the synthesis of phosphatidylserine (PS) (*CHO1*) and linoleic acid (*FAD3* and *FAD2*), which are important components of the cell membrane, were also downregulated. The *cho1*Δ mutant, which is defective in PS synthesis, was shown to exhibit unmasking of beta(1,3)-glucan [63] and is also essential for cell wall integrity [65]. SNP+SHAM treated cells also led to the downregulation of a group of ergosterol biosynthesis pathway genes (Fig 4B). Inhibition of sterol biosynthesis has been shown to lead to an increase in the incidence of lipid droplets [66], which we also observed in SNP+SHAM treated cells. *UPC2*, encoding a transcription factor known to activate ergosterol biosynthesis, especially under hypoxic conditions, was upregulated and required for the observed resistance to caspofungin. Decreased ergosterol production has previously been observed in respiration-deficient *C. albicans* mutants [67,68]. An increase in *UPC2* expression may be a response to reduced ergosterol production, or alternatively SNP+SHAM treatment may mimic hypoxia, a condition that also leads to elevated Upc2. In agreement with this we observed a significant overlap in differentially expressed genes between SNP+SHAM treatment and published microarray data produced under conditions of hypoxia (Fig S6) [69]. The authors suggest that the decrease in ATP-generating activity during hypoxia, which would also occur with SNP+SHAM inhibition, may be an important signal by mitochondria which can result in transcriptional changes. Therefore, respiration inhibition and hypoxia may share common, but as yet undefined, pathways of signalling.

The respiration deficient mutant *goa1*Δ was shown to have an altered cell wall, with an overall decrease in cell wall hexose content, which was attributed to a reduction in high- and intermediate-molecular weight mannans [17]. Shorter outer cell wall fibrils were also seen in *goa1*Δ by TEM, similar to SNP+SHAM treated cells (Fig 2B, C). Mutations that affect mannan composition have also been shown to be important for recognition by macrophages [70]. Disruption of the mannan layer may also serve to expose underlying ligands in the inner wall. However, there were no significant changes in bulk glucan, chitin or mannan levels for the transient inhibition in the case of SNP+SHAM treatment, which acts mainly on Complex IV and the alternative pathway. In addition, the reduced recognition of the *goa1*Δ mutant by macrophages, which is opposite to that seen in SNP+SHAM treated cells, suggests that cell wall changes resulting from transient respiration inhibition are distinct from those seen in the *goa1*Δ mutant.

Despite unmasking leading to an increase in immune cell recognition SNP+SHAM pre-treatment led to enhanced virulence in a mouse model. Exposure of cell wall beta glucan has been shown to increase the inflammation response [71]. One possibility therefore is that SNP+SHAM pre-treatment may have led to an increased and damaging inflammation response. Another possibility rests with the activation of hyphal switching that we observed upon removal of SNP+SHAM, which would have occurred prior to injection. A more rapid hyphal growth response may have facilitated escape from macrophages as well as damage to epithelial barriers, aiding dissemination of the infection. The same rapid transition to hyphal growth was observed when cells were pre-treated with cyanide alone, suggesting that inhibition of the classical ETC is the major cause for this phenomenon. The absence of a rapid hyphal response in the similarly pre-treated *aox2-aox1*Δ mutant suggests that an increased level of Aox2 could be important for the rapid switch to hyphae in response to stress. This observation, together with previous evidence that the alternative pathway is more active in hyphal cells [72], warrants further investigation into the precise role of Aox2 in morphogenetic switching.

In addition to hyphal growth regulators, several other genes previously found to be required for virulence were upregulated by SNP+SHAM treatment. For example Tsa1 (thioredoxin peroxidase 1), which provides resistance against reactive nitrogen species and has also been shown to contribute to fungal virulence [73] was among the most highly upregulated genes following SNP+SHAM treatment. Other upregulated genes associated with pathogenesis included *SAP6* [74], *GLK1*, *PHO112* [75], *PLB1* [76] and the transcription factor *GAT1* as well as its regulation targets *DUR1,2* and *MEP2* [77]. The induction of these genes, several of which are normally associated with the hyphal form of *C. albicans*, suggests that SNP+SHAM treatment may prime cells for increased virulence which could manifest upon withdrawal of the treatment.

In summary our study suggests that the inhibition of respiration as an antifungal strategy in *C. albicans* is a complex proposition. Although it is possible to inhibit respiration with chemicals well tolerated by humans such as SNP+SHAM, this act would appear to lead to changes that may aid and inhibit virulence. On the one hand respiration inhibition clearly leads to unmasking and enhanced recognition by macrophage cells, while on the other primes a stress response leading to increased filamentation and increased virulence. We would argue that the development of a mitochondrial inhibition strategy, while still attractive given the importance of the organelle for *C. albicans* growth, requires a better understanding of the signalling properties of this organelle under conditions of stress.

## Materials and methods

### *C. albicans* growth conditions and chemicals

*C. albicans* strains were maintained on YPD agar plates and grown in YPD in a 30 °C shaking incubator unless stated otherwise. The concentrations of sodium nitroprusside dihydrate (SNP) and salicylhydroxamic acid (SHAM) (Cat. No. 1.06541 and S607, Sigma-Aldrich, Dorset, UK) used for inhibition were 1 mM and 0.5 mM respectively, added to log phase cells followed by 18 h growth unless stated otherwise. Potassium cyanide, Calcofluor White, Congo Red and caspofungin diacetate (Cat. No. SML0425) were obtained from Sigma-Aldrich.

The transcription factor deletion library used in this study was constructed by Homann et al. [32]. Most of the mutants screened, including the *sko1*Δ mutant, were derived from this library unless stated otherwise. The *upc2*Δ mutant was kindly provided by Prof. Joachim Morschhäuser [33]. The *aox2*-*aox1*Δ mutant was constructed from SN87 using the strategy described in [34]. Briefly, *LEU2* and *HIS1* were amplified from plasmids pSN40 and pSN52 respectively, using universal primers with 80 bases homologous to the 5’ end of *AOX2* and the 3’ end of *AOX1* ORF’s. This strategy was designed to delete both *AOX2* and *AOX1* simultaneously as well as the region between these two adjacent genes. The primers used were AOX2-UP2: 5’ atgcttactgcttcgctttacaaacaattaccggtgttaaccaccacagctacttcaacatattctttcattagattatcACCAGTGTGATGGATATCTGC 3’ and AOX1-UP5: 5’ CtaaagatacaaatcctttctttcccatccttggggtctagttacatctaaattgtaatttggttgtggcttgtctgaatAGCTCGGATCCACTAGTAACG 3’. The PCR products were then transformed sequentially into *C. albicans* SN87 using an electroporation protocol [35], followed by selection on agar plates containing Yeast Nitrogen Base without amino acids supplemented with 2% glucose and -His or -Leu dropout powder (Formedium, UK) as appropriate. A summary of strains used in this study is shown in Supplementary Table S1.

### Whole cell respirometry

Respirometry was carried out in real time using an Oxygraph-2k respirometer (Oroboros Instruments, Austria) which was calibrated at 30 °C as per the manufacturer’s instructions. Cells from an overnight culture in YPD were added to 3 ml fresh YPD to a final optical density at 600 nm (OD_600_) of 0.2. The cells were incubated at 30 °C with shaking for 2 h. The cells were then counted and diluted in YPD to give a final cell concentration of 1 ×; 10^6^cells/ml, of which 2.5 ml was added to each chamber in the respirometer. Routine respiration refers to the respiration level immediately prior to the addition of inhibitors. SNP was added to a final concentration of 1 mM, followed by a second addition to a final concentration of 2 mM after 20 min. SHAM was added to a final concentration of 1 mM followed by a second addition 10 min later to give a final concentration of 2 mM. Lastly, potassium cyanide was added to give a final concentration of 2 mM. Data was analysed using Datlab 6 software (Oroboros Instruments). Six independent experiments were performed.

### Viability assays

Cells from an overnight culture in YPD were diluted in YPD to a final OD_600_ of 0.2. The cells were grown for 5 hours at 30 °C with shaking. SNP and SHAM were added to a final concentration of 1 mM and 0.5 mM respectively and cultures were grown for a further 18 h. The cells were collected, washed three times in PBS and counted using a haemocytometer, then diluted in PBS. Four hundred cells of each suspension were plated on YPD agar plates. The plates were incubated for 24 h and the number of colony forming units (CFUs) were counted and compared to untreated controls. Three independent experiments were performed.

### Cell wall agent susceptibility assay

YPD plates were prepared containing either 1 mM SNP, 0.5 mM SHAM, 25 μg/ml Calcofluor White, 50 μg/ml Congo Red, or combinations of these as stated. Cells from an overnight culture in YPD were washed three times in PBS and diluted in PBS to a final OD_600_ of 0.2. Cells were serially diluted and equal volumes were spotted on the plates using a replica plating tool. The plates were incubated for 48 h at 30 °C and photographed.

### Antifungal susceptibility assays

A library of transcription factor mutants [32] was screened using a combination of 100 ng/ml caspofungin, 1 mM SNP and 0.5 mM SHAM in synthetic complete medium, 2 % glucose, 50 mM MOPS pH 6. Each strain was grown overnight in YPD at 30 °C, then washed 3 times in PBS. The OD_600_ was measured and used to inoculate 0.5 ml in 48 well plates to achieve a final OD_600_ of 0.1. Growth of the strains was monitored at 30 °C using a BMG labtech SPECTROstar Nano plate reader with orbital shaking at 400 rpm.

Microdilution assays were carried out in 96-well plates with caspofungin, with or without 1 mM SNP + 0.5 mM SHAM, using synthetic complete medium, 2 % glucose, 50 mM MOPS pH 6. A dilution series of caspofungin was made using the appropriate diluent between 3.9 ng/ml – 8 μg/ml. *C. albicans* cells from an overnight YPD culture were washed three times in PBS. The OD_600_ was measured and adjusted to 2.0. This cell suspension was diluted 1:100 into the appropriate diluent. One hundred microliters of cell suspension was added to the 100 μl in each well of the dilution series. The plates were incubated at 30 °C for 24 h without shaking before determining caspofungin MIC_80_ (the drug concentration needed to decrease the OD_600_ by 80% compared to untreated cells). Three independent experiments were performed.

### RNA isolation and RNAseq

*C. albicans* SC5314 was grown for 5 h in YPD at 30 °C in a shaking incubator. SNP and SHAM were added to final concentrations of 1 mM and 0.5 mM respectively and the cells were returned to the incubator for a further 30 min. RNA was then extracted using a E.Z.N.A.^®^ Yeast RNA Kit (Omega Bio-tek, Norcross, GA) following the manufacturer’s instructions, for three biological replicates per group. RNA was sent to the Centre for Genome Enabled Biology and Medicine (Aberdeen, UK), who performed preparation of stranded TruSeq mRNA libraries, QC/quantification and equimolar pooling, and sequencing on an Illumina NextSeq500 with 1×75bp single reads and average depth of 30M reads per sample. Raw RNAseq data was analysed using the suite of tools available on the Galaxy platform [36]. Briefly, reads were aligned to Assembly 21 of the *C. albicans* genome (Candida Genome Database [37]) using HISAT2. Differentially expressed genes between untreated and treated samples were identified using Cuffdiff. The data discussed in this publication have been deposited in NCBI’s Gene Expression Omnibus and are accessible through GEO Series accession number GSE114531 (https://www.ncbi.nlm.nih.gov/geo/query/acc.cgi?acc=GSE114531).

### *C. albicans* adhesion assay

The adhesion of *C. albicans* to serum-coated polystyrene was assessed. One hundred microlitres of 50% FBS in PBS (Gibco, Thermo Fisher Scientific) was added to the wells of a polystyrene 96-well plate (Corning, USA) and the plate was incubated for 90 min at 37 °C. The wells were then washed three times with PBS. Untreated cells or cells grown in the presence of 1 mM SNP + 0.5 mM SHAM for 18 h were washed three times with PBS and the OD_600_ was measured. The OD_600_ was adjusted to 0.2 in PBS and 100 μl of cell suspension was added to each well. The plate was incubated at 37 °C for 2 h, after which non-adherent cells were removed by washing three times with PBS. The relative biomass of adherent cells was determined using a crystal violet staining-destaining protocol [38].

### Cell wall staining

Cells from an overnight culture in YPD were diluted in YPD to a final OD_600_ of 0.2 and grown for 5 h at 30 °C. 1 mM SNP and 0.5 mM SHAM were added to the cultures and they were grown for a further 18 h. The cells were collected and washed three times in PBS. Cells were then stained with 25 μg/ml Wheat Germ Agglutinin, Alexa Fluor™ 594 Conjugate (Cat. No. W11262, Thermo Fisher Scientific, Waltham, MA) in PBS at room temperature for 1 h in the dark. After washing three times in PBS, the cells were examined using a RFP filter with low-level brightfield illumination with an Olympus IX81 inverted microscope illuminated using a CoolLED pE-4000 unit and captured using an ANDOR Zyla 4.2 CMOS camera. For dectin-1 staining, the cells were incubated with 5 μg/ml Dectin-1-Fc (a kind gift from Prof. Gordon Brown, MRC Centre for Medical Mycology, at the University of Aberdeen) in PBS, 2% BSA for 1 hour at 4 °C. The cells were washed three times in PBS and incubated with sheep anti-human antibody-FITC (Thermo Fisher Scientific, Cat. No. PA5-16924) for 2 h at 4 °C. After washing three times in PBS, the cells were examined by microscopy using a GFP filter with low-level brightfield illumination. WGA and dectin-1 staining of the cell wall were manually assessed using ImageJ v1.50 (NIH, Bethesda, MD). To calculate the percentage of cells with lateral wall staining by WGA, staining of bud scars was ignored. Dectin-1 staining of the cell wall was either scored manually per cell or analysed based on mean fluorescence. To calculate this corrected total cell fluorescence, the background measurement was subtracted from the total integrated density for each whole microscopy image. This result was then divided by the total number of cells in the image. At least 400 cells were counted for each experiment and three independent experiments were performed per analysis.

### Western blotting

Cells from an overnight culture in YPD were diluted in YPD to a final OD_600_ of 0.2. The cells were grown for 5 h at 30 °C with shaking. SNP and SHAM were added to the cultures to a final concentration of 1 mM and 0.5 mM respectively and the cells were incubated for a further 18 h. Samples were taken to obtain cell pellets of 30 mg (fresh weight), after 1 h and after 18 h following drug addition. To monitor Aox2 expression in response to NO, 1 mM SNP was added and samples taken at 10, 20, 30 and 120 min. The cell pellets were snap frozen at −80 °C. Total protein was extracted by homogenisation at 4 °C with glass beads in the presence of 125 mM Tris-HCl, 2% SDS, 2 % glycerol, 0.14 M 2-mercaptoethanol, bromophenol blue buffer at 4 °C. The samples were run on a 5% stacking, 12.5% resolving SDS-polyacrylamide gel. Proteins were transferred to PVDF membrane using a semi-dry transfer system (Bio-Rad, Watford, UK). Phosphorylated Hog1 or Mkc1 were detected using p44/p42 (1:2000) or p38 (1:1000) monclonal antibodies (Cat. No. 4695 and 8690 respectively, Cell Signalling Technology, Netherlands) followed by a goat anti-rabbit-HRP secondary antibody (1:5000) (A0545, Sigma-Aldrich). A monoclonal antibody against *Sauromatum guttatum* Aox which recognises *C. albicans* Aox2 was used for Aox immunoblotting (1:100) (AS10 699, Agrisera, Sweden). Secondary binding of goat anti-mouse-HRP antibody (1:5000) (Sigma-Aldrich A9917) was detected by ECL and images captured using a Syngene GBox Chemi XX6 system. Blots were stripped using Restore™ Western Blot Stripping Buffer (Thermo Fisher Scientific, Cat. No. 21059) and re-probed for actin using an antibody raised against *S. cerevisiae* actin (a kind gift from Prof. John Cooper, University of Washington) at a 1:1000 dilution.

### Cell wall isolation and HPLC

Cell wall material was isolated and acid hydrolysed in 2M trifluoroacetic acid from untreated cells and cells grown in the presence of 1 mM SNP and 0.5 mM SHAM for 18 h in triplicate[39]. To determine the relative amounts of chitin, glucan and mannan, the hydrolysates were analysed by high-performance anion-exchange chromatography [40].

### Phagocytosis assay

J774.1 murine macrophages (a kind gift from the lab of Prof. Carol Munro, Institute of Medical Sciences, University of Aberdeen) were maintained in DMEM with 10% FBS, 200 U/ml penicillin/streptomycin (respectively, Cat No. 10569010, 10082147, 15070063, Gibco, Thermo Fisher) at 37 °C, 5% CO_2_. Cells were counted and diluted in fresh medium to give 5 ×; 10^4^ cells in 0.3 ml then added to the wells of a 8 well μ-Slide (ibidi GmbH, Germany). The cells were then incubated overnight. *C. albicans* was grown in the presence of 1 mM SNP and 0.5 mM SHAM for 18 h. *C. albicans* cells were washed three times in PBS and counted, then diluted to 1.5 ×; 10^4^cells in 0.3 ml in macrophage culture medium with 10 μg/ml Calcofluor White and vortexed briefly. This cell suspension was then added to macrophages. The cells were then co-incubated for 1 h and examined by microscopy. Uptake of *C. albicans* was manually assessed using ImageJ. At least 200 *C. albicans* cells were counted for each experiment.

### Hyphal induction

*C. albicans* grown in the presence of 1mM SNP and 0.5 mM SHAM for 18 h were washed three times in PBS and the OD_600_ was measured. Cells were added to DMEM + 10 % FBS with 10 μg/ml Calcofluor White to a final OD of 0.1 and added to a 8 well μ-Slide (0.3 ml per well). Following a 90 minute incubation at 37 °C, 5% CO_2_, the cells were examined by microscopy using a DAPI filter. The percentage of hyphal cells was determined manually using ImageJ. A total of least 500 cells were counted for each experiment.

### Zebrafish *in vivo* imaging

Two-day old zebrafish of the *Tg(mpeg1:mCherryCAAX)sh378)* transgenic strain were used in this study. Zebrafish were maintained according to standard protocols. Adult fish were maintained on a 14:10 – hour light / dark cycle at 28 ^o^C in UK Home Office approved facilities in the Bateson Centre aquaria at the University of Sheffield. *C. albicans* grown in the presence of 1 mM SNP and 0.5 mM SHAM for 18 h were washed three times in PBS, counted and adjusted to give 500 colony forming units (CFU) in 1 nl and pelleted by centrifugation. Pellets were resuspended in 10% Polyvinylpyrrolidinone (PVP), 0.5% Phenol Red in PBS. The injection and imagining of zebrafish larvae was performed as described in [41] with the exception that the larvae were not immobilised in agar channels and were instead aligned manually in E3 containing 0.168 mg/mL tricaine in glass-bottomed, 96-well plates for imaging. Assessment of *C. albicans* uptake by macrophages was performed manually using NIS-Elements Viewer 4.20 (Nikon, Richmond, UK). Three independent experiments were performed using 10 larvae per condition.

### Electron microscopy

*C. albicans* was grown in the presence of 1mM SNP and 0.5 mM SHAM for 18 h. A 1 ml sample of each culture was centrifuged and the cell pellet frozen under pressure using a Leica EM AFS2 automatic freeze substitution system and EM FSP freeze substitution processor (Leica Microsystems). Samples were dehydrated in anhydrous acetone containing 1% OsO_4_ for 10 h, sequentially warmed to −30°C over 8 h in acetone/OsO4, to 20 °C over 3 h in acetone and then embedded in increasing concentrations of Spurr’s (epoxy) resin over 24 h. Survey sections of 0.5 mm thickness were stained with toluidine blue to verify optimal cell density. All sections were cut with a 45° diamond knife (Diatome US, Hatfield, PA) using a Leica UC6 ultramicrotome. Ultrathin (100 nm) sections were adhered to 300 nm copper mesh grids for examination. Sections were stained with uranyl acetate and lead citrate at 20°C using a Leica AC20 EM automated staining machine. The freeze substitution and embedding, sectioning and staining was carried out by the Microscopy and Histology Facility of the Institute of Medical Sciences, University of Aberdeen. Examination of ultrathin sections was done with a JEOL 1400 Plus transmission microscope (JEOL UK Ltd.) operating at 80 kV and images were recorded using an AMT ActiveVu XR16M camera (Deben UK Ltd.). The AMT v602 camera software was used to visualise, edit and save the images. ImageJ was used to manually measure cell wall thickness. Five measurements were taken along the cell wall for each cell, for a total of 25 cells per group.

### Lipid droplet staining

The lipid droplet stain LD540 was a kind gift from C. Thiele [42]. Wild-type *C. albicans* in log phase was treated with 1 mM SNP and 0.5 mM SHAM for 18 h. A sample of the culture was washed three times in PBS and stained with 0.1 μg/ml LD540 for 30 min at room temperature. The cells were washed three times in PBS and examined using an RFP filter with low-level brightfield illumination.

### Murine *C. albicans* systemic infection model

The effect of SHAM+SNP pre-treatment of *C. albicans* was evaluated using a murine intravenous challenge assay. Pre-treatment of *C. albicans* was performed as follows: cells from an overnight culture in YPD were diluted in YPD to a final OD_600_ of 0.2. The cells were grown for 5 hours at 30 °C with shaking. SNP and SHAM were added to a final concentration of 1 mM and 0.5 mM respectively and cultures were grown for a further 18 h. BALB/c female mice (6-8 weeks old, Envigo UK) were randomly assigned into groups of 6, with group size determined by power analyses using data previously obtained using this infection model. Mice were acclimatized for 5 days prior to the experiment. Mice were weighed and tail-marked using a surgical marker pen to allow for identification. Food and water was provided *ad libitum*. *C. albicans* cells were washed twice with sterile saline and diluted in sterile saline to produce an inoculum of 4 x10^4^ CFU/g mouse body weight in 100 μl PBS. Inoculum level was confirmed by viable plate count on Sabouraud Dextrose agar. Mice were weighed and checked daily until day 3 post-infection when all mice were culled by cervical dislocation. The kidneys were removed aseptically for fungal burden determination, with kidneys weighed and homogenised in 0.5 ml sterile saline. Dilutions were plated on Sabouraud Dextrose agar and incubated overnight at 35 °C. Colonies were counted and expressed as colony forming units (CFU) per g of kidney. Change in weight was calculated as percentage weight change relative to starting weight. An outcome score was calculated based upon kidney burdens and weight loss at time of culling [43].

### Ethics statement

Mouse experiments were carried out under licence PPL70/9027 awarded by the UK Home Office to Dr Donna MacCallum at the University of Aberdeen. All experiments conform to the UK Animals (Scientific Procedures) Act (ASPA) 1986 and EU Directive 2010/63/EU. Zebrafish work was performed following UK law: Animal (Scientific Procedures) Act 1986, under Project License PPL 40/3574 and P1A4A7A5E. Ethical approval was granted by the University of Sheffield Local Ethical Review Panel under project licenses 40/3574 and P1A4A7A5E.

## Acknowledgements

We thank Professor Gordon Brown (MRC Centre for Medical Mycology, University of Aberdeen) for providing Dectin-1-Fc. We would also thank the Microscopy and Histology Core facility Institute of Medical Sciences, University of Aberdeen where the high pressure freezing was performed and staff at the Centre for Genome Enabled Biology and Medicine (University of Aberdeen Medical Research Facility) who performed RNA sequencing procedures. We would also thank members of both the Aberdeen Fungal Group, Kent Fungal Group and WTSA MMFI consortia for comments and useful discussion throughout the project.

## Supplementary Figure legends

**Fig S1. SNP+SHAM treatment does not significantly affect *in vitro* adhesion.**

*C. albicans* wild type and *bcr*1Δ mutant cells were grown with 1 mM SNP + 0.5 mM SHAM for 18 h, then washed and adhesion to serum-coated polystyrene was assessed as described in materials and methods. Graphs show means ± standard deviation. Student’s t-test was used to compare groups, *p<0.05

**Fig S2. SNP+SHAM induces surface exposure of β(1,3)-glucan in both the wild-type and *upc2*Δ**

Wild type strain and *upc2*Δ mutant cells were treated with 1 mM SNP + 0.5 mM SHAM for 18 h and stained with dectin-1 as described in materials and methods. Dectin-1 staining of the cell wall was assessed manually from microscopy images, n=3. Three independent experiments were analysed. Graphs show means ± standard deviation. Student’s t-test was used to compare groups, * p<0.01

**Fig S3. Surface exposure of chitin and β(1,3)-glucan is increased in *ndh51*Δ relative to the wild-type strain.**

Samples from overnight cultures of the wild-type and *ndh51*Δ cells were washed three times in PBS and stained with **(A)** FITC-WGA or **(B)** dectin-1. Staining of the cell wall was scored manually from microscope images, n=3. Three independent experiments were analysed in each case. Graphs show means ± standard deviation. Student’s t-test was used to compare groups, **p<0.01, *p<0.05

**Fig S4. Inhibition classical ETC in *C. albicans* leads to enhanced uptake by murine macrophages**

**(A)** Wild-type and *aox2-aox1*Δ *C. albicans* cells were grown in the presence of 1 mM KCN or 0.5 mM SHAM for 18 h as indicated. The cells were washed and co-incubated with macrophages as described in materials and methods and uptake scored manually from microscope images taken after 1 h, n=3. **(B)** The same procedure was applied to wild type and *ndh51*Δ cell*s*, n=3. Three independent experiments were analysed in each case. Graphs show means ± standard deviation. Student’s t-test was used to compare groups, * p<0.01

**Fig S5. Inhibition of classical ETC with cyanide enhances filamentation upon withdrawal of inhibition**

Wild-type *C. albicans* was grown in the presence of 1 mM KCN in YPD for 18 h. Cells were then washed three times in PBS and transferred to DMEM + 10% FBS at 37 °C. After 90 min incubation, the cells were examined for hyphal growth by microscopy. Representative examples are shown in **(A). (B)** Summary of the percentage of filamentous cells from two independent experiments.

**Fig S6. Comparison of transcriptomes of SNP+SHAM treated cells and cells in early hypoxia.**

Differentially expressed genes induced by SNP+SHAM treatment were compared to those identified within microarray data by Sellam et al. [69] examining the early response to hypoxia, using data from the 30 min time point. Selected genes common to both datasets are highlighted. Up-and down arrows indicate their up-or downregulation respectively.

**Table S1. Strains Used in This Study**

All yeast strains and their origins are listed

**Table S2. List of genes differentially expressed upon SNP treatment**

Cells were treated with 0.5 mM SNP for 30 min and processed for RNA Sequencing and downstream analysis as described in materials and methods. A full list of differentially expressed genes is provided.

**Table S3. List of genes differentially expressed upon SHAM treatment**

Cells were treated with 1.0 mM SHAM for 30 min and processed for RNA Sequencing and downstream analysis as described in materials and methods. A full list of differentially expressed genes is provided.

**Table S4. List of genes differentially expressed upon SNP+ SHAM treatment**

Cells were treated with 0.5 mM SNP and 1.0mM SHAM for 30 min and processed for RNA Sequencing and downstream analysis as described in materials and methods. A full list of differentially expressed genes is provided.

